# A safer, urea-based in situ hybridization method improves detection of gene expression in diverse animal species

**DOI:** 10.1101/133470

**Authors:** Chiara Sinigaglia, Daniel Thiel, Andreas Hejnol, Evelyn Houliston, Lucas Leclère

## Abstract

In situ hybridization is a widely employed technique allowing spatial visualization of gene expression in fixed specimens. It has proven to be essential to our understanding of biological processes, including developmental regulation. In situ protocols are today routine in numerous laboratories, and although details might change, they all include a hybridization step, where specific antisense RNA or DNA probes anneal to the target nucleic acids strand. This step, in general, is carried out at high temperatures and in a denaturing solution, the hybridization buffer, commonly containing 50% (v/v) formamide. An important drawback is that hot formamide poses a significant health risk and so must be handled with great care.

We were prompted to test alternative hybridization solutions for in situ detection of gene expression in the medusa of the hydrozoan *Clytia hemisphaerica,* where traditional protocols caused extensive deterioration of the morphology and texture during hybridization, hindering observation and interpretation of results. Inspired by optimized protocols for Northern and Southern blot analysis, we substituted the 50% formamide with an equal volume of 8 M urea solution in the hybridization buffer. The new protocol yielded better morphologies and consistency of tissues, and also notably improved the resolution of the signal, allowing more precise localization of gene expression, as well as reduced staining at non-specific sites. Given the improved results using a less toxic hybridization solution, we tested the urea protocol on a number of other metazoans: two brachiopod species (*Novocrania anomala* and *Terebratalia transversa*) and the worm *Priapulus caudatus,* obtaining a similar reduction of aspecific probe binding. Overall, substitution of formamide by urea in in situ hybridization offers safer alternative protocols, potentially useful in research, medical and teaching contexts. We encourage other workers to test this approach on their study organisms, and hope that they will also obtain better sample preservation, more precise expression patterns and fewer problems due to aspecific staining, as we report here for *Clytia* medusae and *Novocrania* and *Terebratalia* developing larvae.

## Introduction

In Situ Hybridization (ISH) is a widely employed and powerful technique, allowing localization of specific DNA or RNA strands within cells or tissues. This coupling of genetic and histological information provides a synthetic view of spatial gene expression. Nucleic acids have the fundamental property of pairing to a complementary sequence, which in this case is exogenously synthesized and properly labeled, allowing detection of known target sequences. The technique was developed in the 60s (Pardue and Gall, 1969), and has since then proven invaluable in cell and developmental biology research, as well as in medical diagnostics.

The method has been successfully applied to animals, plants and bacteria, and over the years numerous protocols have been developed, tailored to specific needs, such as the detection of non-coding RNA, or different sample types, including tissue sections, whole mount embryos or cell preparations. The labeling and subsequent detection of probes are variable, and if probes historically incorporated radioactive nucleotides, they nowadays mostly use safer alternatives, such as biotin or Digoxigenin (DIG)-linked nucleotides (Tautz and Pfeifle, 1989), or again fluorescent or enzymatic tags.

The hybridization step, central to the process, is carried out at high temperatures – usually a temperature in the 55° to 65°C range is chosen as a good compromise between sensitivity and specificity - which promote the breaking of hydrogen bonds and destabilize the nucleic acid strands. The ideal temperatures for denaturation and annealing depend on the nature of the target nucleic acids strands, and are usually quite high: denaturation temperature, or melting temperature (Tm) is determined by the sequence’s base composition (C-G Watson-Crick bonds are more stable than A-T), with ideal hybridization temperatures about 25°C below Tm (Marmur and Doty, 1961). Unfortunately, the long incubations usually performed increase the risk of nucleic acid degradation and tissue damage in the samples, and therefore much effort has been dedicated in the past towards the best method to lowering hybridization temperatures. Early reports used factors such as salt concentration, pH, and solvents to modulate the efficiency and stringency of the hybridization process, and, more importantly, to favor destabilization of DNA or RNA chains, thus lowering the reaction temperature. Various organic solvents were found to reduce the stability of nucleic acid strains, including guanidinium chloride, salicylate, formamide, dimethyl sulfoxide (DMSO), N,N’-dimethylformamide (DMF), a variety of alcohols (for example see (Rice and Doty, 1957; Marmur and Ts’o, 1961; Hamaguchi and Geiduschek, 1962; Herskovits, 1962; Levine et al., 1963)), urea and several of its derivatives, or also sodium hydroxide, used in the first hybridization in situ on *Xenopus* oocytes (Pardue and Gall, 1969).

Initial reports of in situ hybridization achieved denaturation either with high temperatures or chemically, with NaOH or salts (John et al., 1969; Buongiorno-Nardelli and Amaldi, 1970; Barsacchi and Gall, 1972; Gall, 2016). Subsequent research favored the use of formamide at a concentration of 50-70 % in the hybridization buffer, in order to lower hybridization temperature and efficiently denature DNA/RNA (see for example (Barbera et al., 1979; Bauman et al., 1980; Gerhard et al., 1981; Hafen et al., 1983; Levine et al., 1983; Braissant and Wahli, 1998; Brown, 1998). This organic solvent was found to be particularly useful, for its ability of denaturing and renaturing DNA at room temperature (Hutton, 1977; Marmur and Ts’o, 1961; McConaughy et al., 1969), a property that allowed the generation of the first DNA-RNA hybrids (Bonner et al., 1967). Formamide is nowadays standardly employed in different hybridization methods, either in situ hybridization or Northern and Southern blotting, and provides generally reliable results. Unfortunately, it is also a very hazardous chemical, causing both short term effects such as respiratory tract irritation, headache and nausea, and long term damages at the level of internal organs and on reproduction ((Fail et al., 1998; George et al., 2002, 2000; Gleich, 1974; Stula and Krauss, 1977; Merkle and Zeller, 1980; Kennedy and Short, 1986), see also Table 1 for further details). It is rapidly absorbed orally, via inhalation or skin contact, and, given that in experimental animals it has been shown to have embryotoxic and teratogenic effects (Merkle and Zeller, 1980; George et al., 2000, 2002), pregnant women are considered to be particularly at risk (*European Chemical Agency. Proposal for identification of a substance as a CMR CAT 1A or 1B, PBT, vPvB or a substance of an equivalent level of concern. Formamide*). Moreover, the generally high temperatures of reaction, pose an additional threat, since augmented evaporation increases the risk of inhalation. These hazards mean that the handling of samples and of waste has to be carefully controlled and managed (*CICAD 31: N,N-Dimethylformamide*, 2001).

**Table 1:**
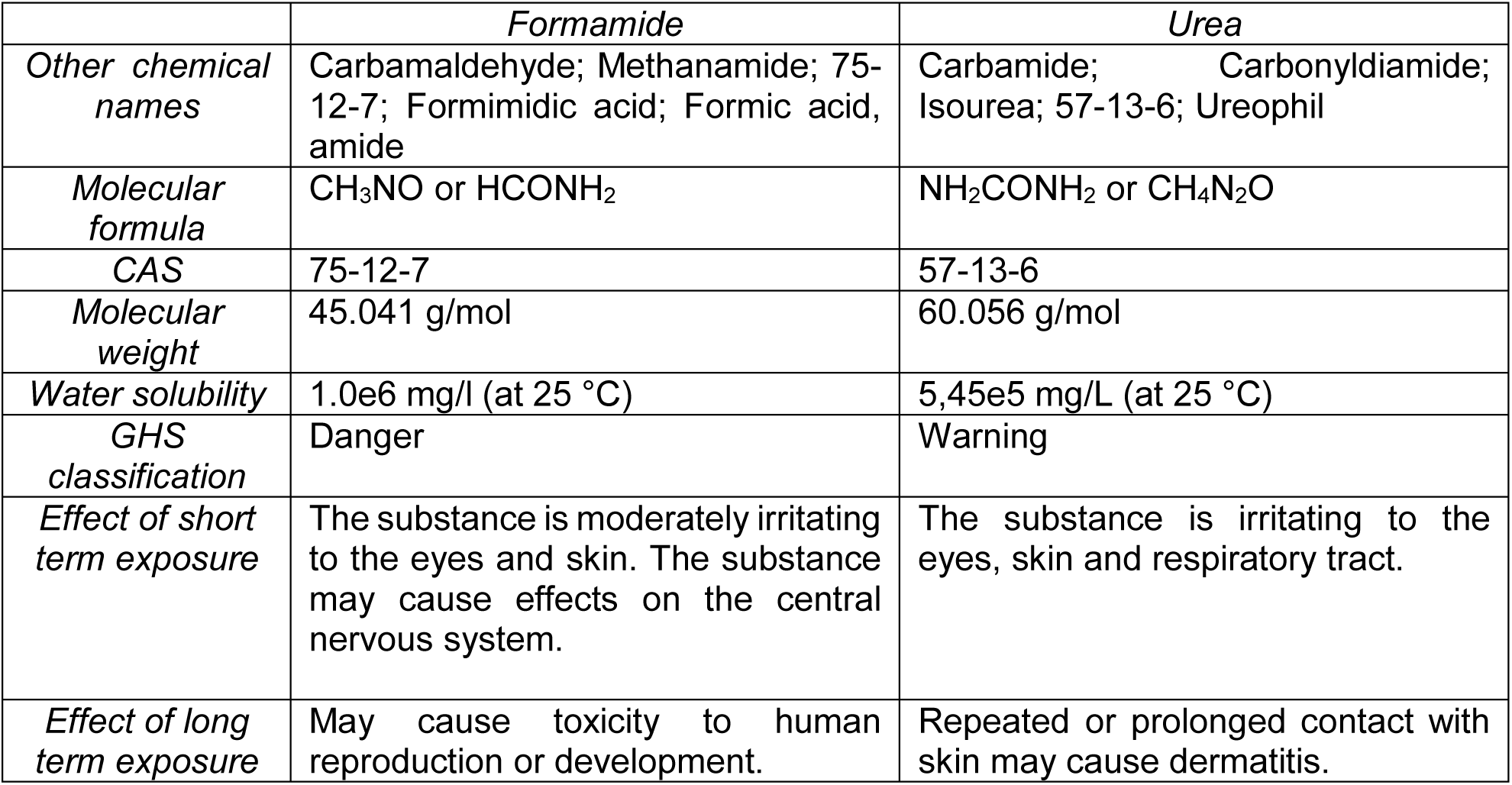
Overview of properties and health risks of formamide, the most diffuse denaturing agent, and urea. Urea represents the safest alternative (source www.ncbi.nlm.nih.gov/pccompound).

The in situ hybridization technique is a well-established method for RNA detection in the hydrozoan *Clytia hemisphaerica*, and has proven valuable for assessing gene expression during embryonic development, oogenesis, and in adult structures such as in the tentacle bulb (Chevalier et al., 2006; Denker et al., 2008; Leclère et al., 2012; Lapébie et al., 2014; Kraus et al., 2015). However, the prolonged, high-temperature hybridization step is rather aggressive for the medusa form, particularly for the fragile umbrella, rich in extracellular matrix. While other morphological features of the medusa, such as the feeding manubrium, the gonads and the tentacle bulbs (see Fig.1A), retain their overall structural integrity, the umbrella becomes deformed and shrunken (Fig.1Bb), thus impairing the study of more fine elements, such as the nervous system network underlying the umbrellar epithelia. This limitation, which different fixation methods could not overcome, prompted us to question the standard hybridization step. We resolved to look into the hybridization buffer composition, searching for alternatives that could improve sample preservation and, since the most abundant reagent was formamide, we decided to target it.

During the 60s urea was identified as another efficient organic solvent (Herskovits, 1963), which is still mainly employed as a denaturing agent in PAGE (Polyacrylamide Gel Electrophoresis) methods (Summer et al., 2009). Indeed, urea and formamide share similar properties, and have been successfully employed for years (Kourilsky et al., 1971) as equivalents in a number of techniques, including Fluorescent In Situ Hybridization (FISH) on bacteria (for a recent report see (Fontenete et al., 2016)), protein denaturation (e.g. (Lim et al., 2009)), or as clearing agents for tissue imaging (e.g. Sca*le*S and ClearT methods, respectively, reviewed in (Azaripour et al., 2016)).

Here, we present a formamide-free in situ hybridization protocol for the hydrozoan *Clytia hemisphaerica*, in which the use of urea as a denaturing agent not only improves the overall morphology of specimens, but can also improve the sensitivity of the detection. Importantly, this substitution provides for a safer, easier procedure, with reduced risks both for the operator and the environment. In addition, we show that this alternative urea-containing hybridization buffer can represent a useful option for in situ hybridizations also in other metazoan species: the protocol was successfully employed for assessing gene expression during development in two brachiopods, *Novocrania anomala* and *Terebratalia transversa*, and in the worm *Priapulus caudatus*.

## Methods

### Animal culture/collection

The *Clytia hemisphaerica* (Linnaeus, 1767) Z4B strain used in this study is cultured in artificial sea water, under controlled conditions of temperature (20°C), pH and water flow, in our in-house aquarium system (Houliston et al., 2010). Medusae were fed with newly hatched *Artemia* and grown until fully mature (3 weeks from release from polyp) before fixation.

*Priapulus caudatus* (Lamarck, 1816) collection was performed as described in (Martín-Durán and Hejnol, 2015), while collection of *Terebratalia transversa* (Sowerby, 1846) and *Novocrania anomala* (O. F. Müller, 1776) was done as in (Santagata et al., 2012) and (Martín-Durán et al., 2016), respectively.

### *Clytia hemisphaerica* in situ hybridization protocol

The protocol was adapted from (Lapébie et al., 2014) and from Takeda et al. (*in preparation*) with regard to the chromogenic (CISH) and the fluorescent in situ hybridization (FISH), respectively. Medusae were relaxed and fixed on ice with a pre-chilled solution of 3.7 % formaldehyde plus 0.4 % glutaraldehyde in 1X PBS (Phosphate-Buffered Saline), for two hours (CISH fixation) or fixed for 36 hours at 18°C with 3.7% formaldehyde in HEM buffer (0.1M HEPES pH 6.9, 50mM EGTA pH 7.2, 10mM MgSO_4_). Specimens were washed thoroughly with 1X PBST (1x PBS plus 0.1% Tween-20), and stepwise dehydrated to 100% methanol, and stored at −20°C. Samples were re-hydrated for 15’ with 50% methanol/ PBST, and then with three PBST washes.

In the hybridization solution, formamide (v/v) (Fig. 1Ba, Ca, Da, Ea, Fa, Ga, Ha, Ia, J, Ka, La, Ma) was substituted with a freshly prepared 8M urea solution (Fig. 1Bb, Cb, Db, Eb, Fb, Gb, Hb, Ib, Kb, Lb, Mb). The final hybridization mix therefore contained: 4M urea, 5X SSC (Saline Sodium Citrate, a buffer solution at pH 7.00), 1% dextran powder (which acts as a volume-excluding polymer to concentrate the probe and increase hybridization rate), tRNA (a blocking agent, reducing non-specific binding), heparin (which reduces background staining (Singh and Jones, 1984)), 1% SDS (Sodium Dodecyl Sulfate, a detergent permeabilizing membranes (Shain and Zuber, 1996)), and milliQ H_2_O to volume. The optimal concentration for the urea solution was determined on the basis of previous reports. (Simard et al., 2001) demonstrated that for Northern blot a 2-4 M urea-containing solution provided the best signal, while a higher concentration significantly decreased the sensitivity of the hybridization. Similarly, (Søe et al., 2011) showed that a 4M urea-containing hybridization buffer provided the best detection of low-copy miRNAs in mouse brains. Samples were gradually transferred to the hybridization solution, and pre-hybridized at 58°C for two hours. Probes were then added at a concentration of 0.1-1 ng, and hybridized at 58°C for 48-72 hours. Samples were then transferred to progressively stringent washes, at 58°C, as follows: 3 X 30 minutes with (4M urea, 0.1% Tween, 5X SSC, milliQ H_2_O), likewise with (2M urea, 0.1% Tween, 5X SSC, milliQ H_2_O), and finally twice for 30 minutes with (0.1% Tween-20, 5X SSC, milliQ H_2_O).

**Figure 1:**
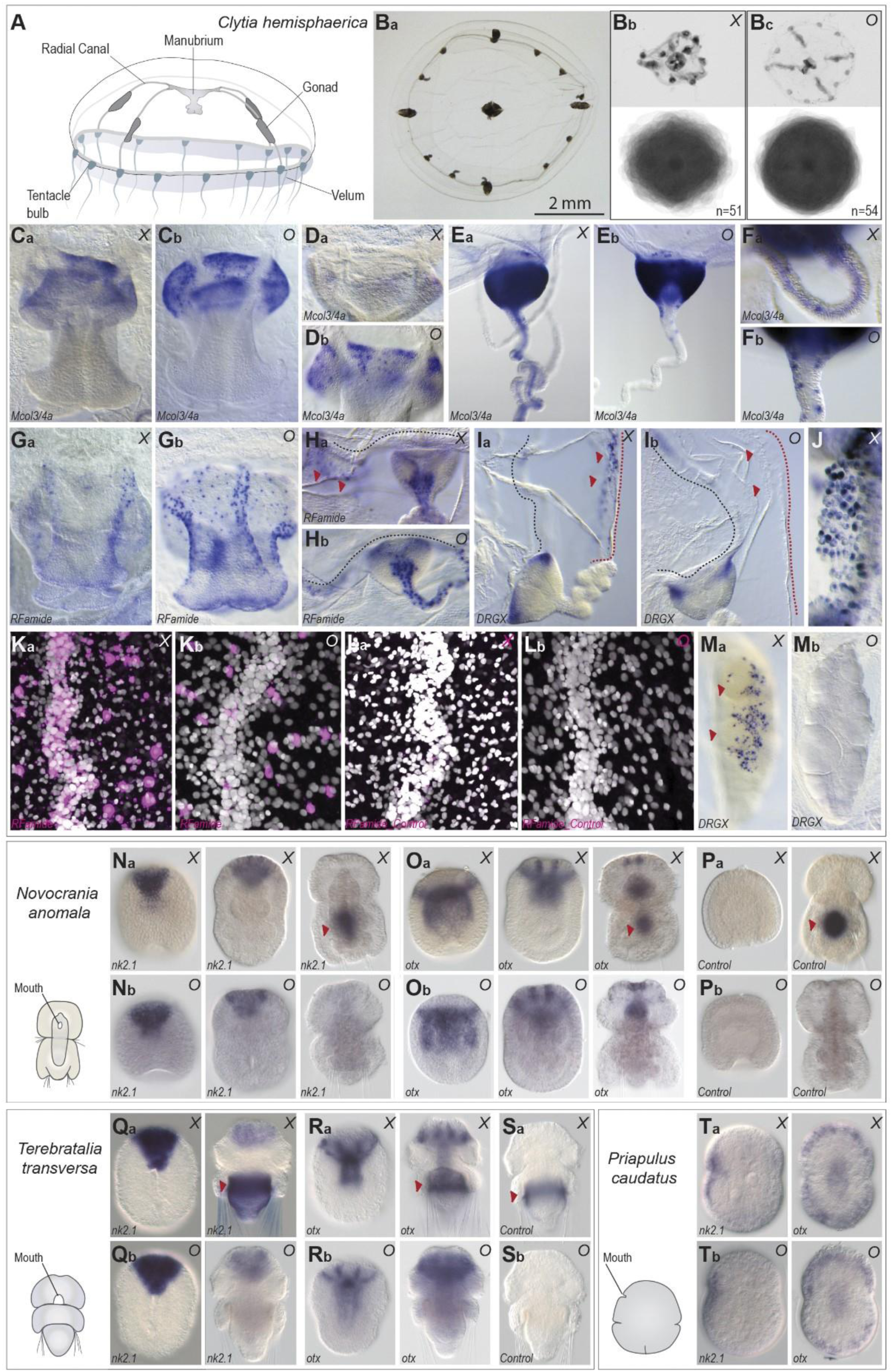
*Comparison of results for* in situ *hybridization with the standard, formamide-based (X), protocol, and the urea-based (O) alternative* **A.** Anatomy of *Clytia* medusa: the tetraradial symmetry is evident in the four radial canals crossing the umbrella, from which the four gonads develop. An acellular, thick, mesoglea separates exumbrella and subumbrella layers. **Ba**. The fixation process efficiently preserves morphology and size of the living medusa. **Bb.** The medusae appear heavily damaged after the formamide-based hybridization step, shape becomes irregular. **Bc.** 4M urea in the hybridization buffer improves overall morphology, and, even if the extensive shrinking of the medusa cannot be avoided, the texture results more flexible. **Ca**. Expression of *minicollagen 3/4a*, in nematoblasts at the base of the manubrium, detected with the formamide protocol. **Cb.** The urea protocol produces a sharper signal at the base of the manubrium. **Da and Db.** With low *minicollagen 3/4a* probe concentration, the manubrium signal is lost with the formamide (Da), while it is still detectable with the urea method. **Ea and Eb.** The resolution of *minicollagen* expression pattern in the tentacle bulb is improved (e.g. single positive cells are visible above the bulb) with the urea-based hybridization buffer (Eb), while with the formamide protocol a non-specific staining in the endoderm of the bulb is frequently observed. **Fa and Fb.** Higher magnification of the *Mcol3/4a* expression in the tentacle of formamide (Fa) and urea (Fb) treated specimens, with the latter showing no endodermal background and improved cellular resolution. **Ga and Gb.** *RFamide* positive cells in the manubrium of a formamide-treated medusa (Ga), however the urea method shows a more complex *RFamide* network (Gb). **Ha and Hb.** *RFamide* positive cells in the tentacle bulb and in the circular canals appear more clearly with the urea method (Gb). In the formamide-based protocol, frequent non-specific staining (red arrowheads) is observed in the velum (Ga). **Ia and Ib.** *DRGX* expression with the formamide (Ia) and urea (Ib) based protocols: aspecific signal (red arrowheads) is seen at the margin of the velum (red dashed line) in Ia. **J.** High magnification of the typical aspecific, superficial signal on the velum obtained with the formamide-based method. **Ka and Kb.** Fluorescent in situ hybridization for *RFamide* with, respectively, formamide-based and urea-based hybridization buffer, a portion of one radial canal and nearby subumbrella is shown. The nuclear staining (DAPI) show that the tissues maintain the overall structure, even if heavily shrunk. Images were taken with the same settings. In Ka the *RFamide* (magenta) signal is shadowed by background staining, while in Kb details are easily distinguishable. **La and Lb**. Sense probe control for *RFamide* with, respectively, formamide and urea methods. Some background staining is seen in the canal of the formamide-treated specimen. **Ma and Mb.** In situ hybridization for *DRGX*, showing the typical non-specific superficial sustaining appearing on the gonads of formamide-treated medusae (Ma). **Na and Nb.** *Nk2.1* expression pattern in formamide (Na) and urea (Nb) –treated embryos (early gastrula, late gastrula and late larva) of the brachiopod *Novocrania anomala*. In the late stage larva, an aspecific signal is seen in the shell gland with the traditional formamide buffer (Na). **Oa and Ob**. *otx* expression pattern in formamide (Oa) and urea (Ob) –treated embryos, same stages as before. Again, the late stage larva shows an aspecific signal in the shell gland, only with the formamide protocol. **Pa and Pb**. The *otx* sense control demonstrates the aspecifity of the shell gland signal. **Qa, Qb, Ra, Rb**. Another brachiopod species, *Terebratalia transversa*, shows a similar pattern of aspecific staining, appearing in the shell glands of late stage larvae when hybridized with formamide (Qa, Ra). **Sa and Sb.** The *otx* sense control demonstrates the non-specificity of signal in the shell glands. **Ta and Tb.** Similar results are obtained with formamide (Ta) or urea (Tb)-based methods, in the case of the worm *Priapulus caudatus*.

For CISH detection, samples were transferred to MABT (Maleic Acid Buffer, containing Tween-20) and incubated with the appropriate antibody (anti-DIG-AP, 1: 2000) overnight at 4°C. They were then transferred to NTMT (NaCl, Tris-HCl at pH 9.5, MgCl_2_, Tween-20). The signal was detected with NBT/BCIP reaction, and then stopped with a rapid milliQ H_2_0 wash, followed by 1x PBS wash. Samples were post-fixed with 3.7% formaldehyde in PBS for 30 minutes, rinsed with 1X PBS, and transferred to glycerol for imaging and long-term storage.

For FISH detection, samples were transferred to MABT, and incubated overnight with the appropriate antibody, peroxidase conjugated (anti-DIG-HRP, 1:2000). Samples were then washed twice in fresh color reaction buffer (0.0015% H_2_0_2_ in PBS) for 30 minutes. Signal was developed with fluorophore-conjugated tyramide kit (Perkin Elmer, 1:400 in color reaction buffer) for one hour. Samples were washed with 1X PBS, stained with Hoechst (1:2000 in 1X PBS) for 30 minutes, rinsed with PBS, and transferred to Citifluor for storage and imaging.

### *Novocrania, Terebratalia* and *Priapulus* in situ hybridization protocols

The protocol was adapted from (Hejnol, 2008), with the following modifications: proteinase K digestion (before hybridization) was followed by a post-fixation with 3.7% formaldehyde and 0.2% glutaraldehyde in PBST, the hybridization solution contained 1% dextran, and, finally, the formamide in the hybridization buffer and stringent-wash buffer, was replaced by 8M urea (with a final concentration of 4M urea). The complete protocol is provided in the supplementary material.

### Image acquisition and processing

Full-sized medusae images were taken on Leica M205 FA and M165 FC stereomicroscopes. Colorimetric images were taken on a Zeiss Axio Imager 2. All fluorescent images were taken on a SP8 Leica confocal microscope, using the same image acquisition and laser parameters. The composite images of medusae (Fig. 1Bb and Bc) were obtained with Photoshop CS5, as follows: images were converted to black-and-white, transformed into outlines with the stylize filter, opacity was reduced to 20%, and the resulting images were overlapped, generating overall shapes. Colorimetric ISH images were adjusted with Photoshop CS5, while fluorescent images were processed with the same noise reduction parameters in the Las X (Leica) software. *Novocrania, Terebratalia and Priapulus* embryos were imaged with Axiocam at a Axioscope 2 using the Axiovision software.

## Results

### Improved medusa morphology following in situ hybridization

The medusa stage of the hydrozoan *Clytia hemisphaerica* displays tetraradial symmetry, with four radial canals running from the oral manubrium to the tentacle bulbs, and four gonads developing around them. The transparent umbrella is composed of an exumbrella layer and a subumbrella, separated by a thick layer of acellular mesoglea. The central manubrium, leading to the digestive pouch, is rather short and also shows tetraradial symmetry. A circular canal runs around the periphery of the umbrella, connecting the 4 radial canals and the tentacle bulbs, numbering 16 in the adult animal. A thin velum, typical of hydrozoan medusae, extends from the umbrella margin and contributes to swimming (Leclère et al., 2016). The fixation process reliably preserved the shape and the size of the live animal, which at the adult stage measures about 1 cm in diameter (Fig.1Ba). The standard, formamide-based, in situ hybridization treatment caused extensive shrinking of the umbrella and altering of the body proportions. The medusae appeared folded and heavily shrunk, while the more conspicuous elements, such as the manubrium, the gonads and the tentacle bulbs, did not appear to be significantly affected, and ultimately formed a scaffold preserving the appearance of the medusa body (Fig. 1Bb). The superposition of images from 51 animals highlights their deformed morphology (Fig. 1Bb). This deformation, along with a newly acquired rather rigid consistency, impaired the analysis of gene expression and the recognition of fine features. On the other hand, the morphology of the medusae following the urea-based treatment was better preserved – as shown by the superposition of 54 different jellyfish (Fig. 1Bc). Although the extensive shrinking could not be overcome, the more flexible consistency under the urea-based protocol, coupled to an improved preservation of the umbrella tissues, greatly facilitated observation of the specimens.

### A more sensitive technique for gene expression analyses

Nervous system-related genes provide a reliable way to assess the precision of mRNA detection, given their cell type specific expression. We thus chose to compare the expression patterns for *minicollagen 3/4a* (Denker et al., 2008), *RFamide,* and *DRGX* (Kraus et al., 2015). In all cases, formamide and urea variants gave comparable expression patterns, demonstrating that 4M urea could efficiently substitute for the 50% formamide during hybridization steps. Interestingly, the urea-based protocol produced sharper staining patterns, particularly in the case of isolated cells, as for example showed by the *Mcol3/4a* expression at the base of the manubrium, and in the tentacle bulbs (Fig. 1C, E, F), where single positive cells could be more easily distinguished.

Additionally, the urea method, with respect to the genes tested, proved to be overall more sensitive than the formamide-based one. This could be seen using highly reduced probe concentrations, when *Mcol3/4a* signal could no longer be detected in the manubrium using the formamide method (Fig. 1Da), while clear staining could still be obtained using urea (Fig. 1Db). In the case of *RFamide* expression, the sensitivity proved even higher, and, using the same probe and color development conditions, the image obtained with the urea method revealed a more complex network of neurons in the manubrium, in the circular canal, and in the subumbrella (Fig. 1G, H).

### An increased signal-to-noise ratio

Side-by-side comparison of staining patterns produced by formamide vs urea protocols also highlighted the tendency of the formamide-based hybridization method to favor aspecific signals. In the case of fluorescent in situ hybridization, which is a highly sensitive method, the background signal appearing with the formamide treatment could be so strong to conceal the true signal, in this case of *RFamide* neurons (Fig. 1La, Lb). Additionally, using a probe against *DRGX* in the CISH method, medusae showed a superficial, punctate, non-specific staining at the margin of the velum (Fig. 1Ib, J) and on the gonads (Fig. 1Mb). The non-specific signal was recognizable because of its fast development time and the superficial localization on the cell layers. These aspecific signals caused premature arrest of the detection reaction and therefore under-development of the ‘true’ expression patterns. Those aspecific staining were not observed using the urea method (Fig. 1Ia, Ma), allowing the development reaction to be continued until detailed expression patterns were obtained.

### The urea method is a reliable alternative for multiple species

The modified, urea-based, in situ hybridization protocol detailed in this study is simple and safe to implement, and could be advantageous for research in other metazoan species.

We thus tested it on different animals and on different developmental stages, posing various experimental challenges. We included two Brachiopoda species *(Novocrania anomala* and *Terebratalia transversa*) and a priapulid worm (*Priapulus caudatus*), for which in situ hybridization protocols have been successfully established, but for which gene expression analysis were at times complicated by aspecific staining. Similarly to *Clytia*, we used the already published nervous system-related genes, *nk2.1* and *otx,* as a reference (Martín-Durán et al., 2016, 2012; Martín-Durán and Hejnol, 2015).

In all three species, the urea-based hybridization buffer produced a specific signal, similar to the one obtained with the formamide-based hybridization buffer (Fig. 1Na-Tb). In the case of *P. caudatus*, the resolution appeared substantially equivalent (Fig. 1Ta, Tb). However, a marked improvement was obtained with the older stages of the two brachiopod larvae, where the aspecific staining that can occur at the shell secreting glands with the formamide hybridization protocol (Fig. 1Na, Oa, Qa, Ra) was absent (Fig. 1 Nb, Ob, Qb, Rb). The aspecific nature of the glandular staining was demonstrated by the signal seen in the corresponding sense-probe formamide control (Fig. 1Pa and Sa), absent in the urea-control.

These results demonstrate that the urea protocol for in situ hybridization can be successfully applied to other species. Furthermore, it can prove useful to prevent probe trapping associated to determined structures of stages, reducing non-specific signal.

## Discussion

The urea-based in situ hybridization protocol described in this study represents an efficient, non-toxic alternative to standard formamide-based in situ hybridization techniques. In both the hydrozoan medusa *Clytia hemisphaerica* and two brachiopod species, *Novocrania anomala* and *Terebratalia transversa*, the urea-based hybridization buffer effectively improved signal detection, reduced aspecific staining and improved specimen morphology.

The common in situ hybridization techniques, routinely employed to reliably assess gene expression in numerous metazoan models, such as *Xenopus*, *Danio*, *Nematostella* or *Mus*, employ large quantities of formamide, a dangerous chemical (see Table 1 for an overview of toxicity of formamide, and its possible substitute urea). The hazard posed by the toxicity of formamide constitutes a problem for its use in the laboratory, however it is usually considered as a necessary step. A typical hybridization buffer contains a 50% volume of formamide, meaning that extreme care is needed both in manipulation and waste disposal. Compounding the danger, the hybridization reaction is carried out at high temperatures (55°C-65°C), posing an additional risk of exposure to vapors due to the increased evaporation. For these reasons, several reports have questioned the extensive use of formamide, asking if a less toxic option, both for health and environmental reasons, could be found (summarized in Table 2). These studies were mostly aimed at medically oriented research, where fluorescent in situ hybridization is a common diagnostic technique for pathogens.

**Table 2:** A survey of the recent reports focused on formamide-alternatives for in situ hybridization. Hybridization types: CISH-Chromogenic In Situ Hybridization, FISH-Fluorescent In Situ Hybridization, FIVH-Fluorescent In Vivo Hybridization, GISH-Genomic In Situ Hybridization.

| *Hybridization type* | *Formamide-substitute* | *Sample type* | *Reference* |
| --- | --- | --- | --- |
| CISH | 4x SSC | Mammal tissue (Fibromatosis nodules) | (Berndt et al., 1996) |
| FISH | Urea-NaCl | Bacteria (*Staphylococcus aureus*) | (Lawson et al., 2012) |
| CISH / FISH | 4M urea | Mammal tissue (miRNA in mouse brain) | (Søe et al., 2011) |
| IQFISH (1 hour-hybridization FISH) | Ethylene carbonate, sulfolane, propylene carbonate, c-butyrolactone, 2-pyrrolidone, d-valerolactam. | Mammal tissue (Breast carcinoma, tonsil and colon tissue) | (Matthiesen and Hansen, 2012) |
| FIVH | 4M urea | Bacteria (*Helicobacter pylori*) | (Fontenete et al., 2013) |
| GISH | 0.02x SSC | Plants (*Prospero, Melampodium*) | (Jang and Weiss-Schneeweiss, 2015) |
| FIVH | 0.5M urea | Bacteria (*Helicobacter pylori*) | (Fontenete et al., 2015) |
| FISH | 4M urea | Bacteria (*Helicobacter pylori*) | (Fontenete et al., 2016) |
| FISH | NaCl-EtOH |  | Sahlgrenska University Hospital (Sweden)<br>www.subsport.eu |
| FISH | 10% dextran sulfate/20% glycerol/0.9% NaCl or KCl |  | Patent WO 1996031626 A1 |
| ISH on paraffin-embedded sections | Chaotropic agents (selected from the group of urea, salts of guanidinium or guanidine) |  | Patent EP 2563935 A1 |

In other types of gene expression analysis, such as Northern, Southern or Western blots, urea is widely employed as a denaturing agent for both proteins and nucleic acids. Indeed, the denaturing properties of urea are known and studied since the 1960s, and the efficiencies of urea and formamide for hybridization to RNA probes on blotting applications have thoroughly been tested (Simard et al., 2001). That study not only demonstrated that urea could effectively replace formamide in detecting RNA, but also found that the best results were obtained when the urea concentration ranged between 2 and 4 M. Above 4M, the sensitivity of detection was markedly reduced, probably due to an increased viscosity of the solution (Hutton, 1977). Our observations confirmed this observation, with the urea-based hybridization solution being more viscous than the formamide one.

In the hydrozoan *Clytia hemisphaerica,* our motivation to search for alternatives to the formamide-based, standard, hybridization step initially lied in the unsatisfactory results obtained on adult medusae, whose texture and morphology were distorted to a point where analyses of gene expression were impaired. In addition, a characteristic array of non-specific staining patterns were frequently observed on the gonads, in the endoderm of the tentacle bulbs, and on the thin velum, casting doubts on the interpretation of newly assessed gene expression patterns. This was particularly problematic when a “salt-and-pepper” distribution was expected -as would be in the case of genes related to nervous system. These issues were greatly improved by substituting the formamide in the hybridization solution and in the stringent washes with an equivalent volume of 8 M urea solution.

Formamide and urea have similar denaturing properties, nevertheless their respective mechanisms of action are still incompletely understood, with multiple factors affecting the efficiency of the reaction. Formamide lowers the melting temperature of DNAs by 2.4 −2.9°C/ mole of formamide - with an efficiency depending on the properties of the nucleic acid strands themselves, such as their G+C content, the helix topology and the state of hydration (Blake and Delcourt, 1996). The solvent weakens hydrogen bonds, ultimately allowing lower hybridization temperatures, with similar high stringencies (Casey and Davidson, 1977; Sadhu et al., 1984; Robertson and Vora, 2012). Generally, the more concentrated the formamide, the higher is the stringency of reaction, but it was shown that, similarly to urea, an excess of solvent causes a dramatic drop in probe binding and signal detection (Manz et al., 1992; Bond and Banfield, 2001). Non-specific signal is a common artifact, and stringency can be further improved through post-hybridization washes, which remove the excess probe and disrupt the incorrectly paired duplexes which might occur – for example, evidence suggests that G and U in RNA can form a weak base pair (Uhlenbeck et al., 1971; Lomant and Fresco, 1975). At this step, the stringency of the washing buffer can also be controlled by lowering the concentration of salt, instead of using formamide, thus reducing the volume of toxic waste (Lathe, 1985). It is worth noticing that another parameter which could affect the signal-to-noise ratio is the purity of formamide, and deionized formamide is recommended in numerous protocols, even if no difference was observed in the case of *Clytia hemisphaerica*, where recently opened bottles of formamide, kept for several weeks at 4°C, are usually employed. The reason for choosing a deionized solvent is that formamide solutions become acidic with time, due to the hydrolytic breakdown of formamide to formic acid and ammonium formate (Chow and Broker, 1989), which attack the phosphodiester bonds of RNA strands, affecting in particular larger RNA molecules. The purification removes the breakdown products, which will then take time to reform, at the reaction conditions.

Due to the widespread use of formamide, few studies have addressed the mechanism of action of urea. Urea can substantially lower the melting temperature of DNA, with values approaching 2°C reduction per mole of urea (Hutton, 1977), thus slightly lower than the decrease that can be obtained with formamide. Recently it was shown that, as hypothesized previously, urea can interact with water and with both polar and nonpolar components of nucleotides - it forms multiple hydrogen bonding with the RNA bases, and generates stacking interactions with them– causing RNA destabilization through a disruption of the base-pair interactions (Herskovits and Bowen, 1974; Priyakumar et al., 2009; Lambert and Draper, 2012). Similarly to our observations, an increase of sensitivity with urea compared to formamide was observed in the case of a FISH protocol developed for detecting *Helicobacter pylori ex vivo,* in gastric biopsies (Fontenete et al., 2013). This might be due to an additional permeabilization role of urea (Lim et al., 2009; Huang et al., 2011), which could enhance probe penetration in the tissues. A permeabilizing activity of urea could explain, for example, the improved detection we obtained in urea-treated medusae with very low probe concentrations, for example for *Mcol3/4* in the manubrium (Fig. 1D).

The urea alternative appears to be a reliable option for routine in situ hybridization, and it has been successfully applied to multiple species, including *C. hemisphaerica*, *N. anomala*, *T. transversa*, *P. caudatus* (this study), the scyphozoan jellyfish *Aurelia aurita* (M. Manuel and T. Condamine, personal communication) and the acoel *Hofstenia miamia* (L. Ricci, personal communication). However, since a previous study on bacteria (Fontenete et al., 2016) reported a certain dependency of sensitivity on the nature of the probe, we would recommend performing an initial comparison, in order to verify the reproducibility of the gene expression patterns detected.

Overall, substitution of formamide by urea in situ hybridization offers a safer alternative protocol, potentially useful in research, medical and teaching contexts. We encourage other workers to test this approach on their study organisms, and hope that they will also obtain more informative and beautiful expression patterns, as we have done in *Clytia hemisphaerica*, *Novocrania anomala* and *Terebratalia transversa.*

## Acknowledgements

We are grateful to Sandra Chevalier for suggesting to focus on the hybridization buffer, Gonzalo Quiroga-Artigas for thorough testing of the urea-protocol, Antonella Ruggiero for troubleshooting of the fluorescent in situ protocol, Stefania Castagnetti and Michael Schubert for kindly sharing their stereoscopes. Research at the Observatoire Océanologique de Villefranche-sur-mer was supported by the ANR grant ANR-13-PDOC-0016 “MEDUSEVO”. Research at the Sars International Centre was supported by the FP7-PEOPLE-2012-ITN grant no. 317172 “NEPTUNE”.

